# Homology-mediated transformation of frog-killing fungus *Batrachochytrium dendrobatidis* illuminates chytrid development and pathogenesis

**DOI:** 10.1101/2025.08.29.673073

**Authors:** Stephanie M. Brody, Erik Kalinka, Sarah Prostak, Tamilie Carvalho, Jarrett Man, Timothy Y. James, Lillian K. Fritz-Laylin

## Abstract

The chytrid fungus *Batrachochytrium dendrobatidis* (*Bd*) infects amphibians and causes chytridiomycosis, a disease linked to global amphibian decline. Despite its ecological importance, *Bd* has lacked robust tools for genetic manipulation, limiting molecular insights into its development and pathogenicity. Here, we establish homologous recombination as a system for stable transformation of *Bd*, enabling targeted chromosomal integration of exogenous DNA, endogenous protein tagging, and targeted gene deletion. We use this system to visualize *Bd* infection in live amphibians to enhance understanding of host invasion and pathogenesis. We also use this system to test a previous hypothesis regarding the role of chitin synthases in *Bd* development by tagging the endogenous chitin synthase Myo17D and observing its rapid relocalization to the plasma membrane during *de novo* cell wall assembly. Finally, we use our homologous recombination approach for targeted gene deletion by knocking out the *URA3* locus, and confirm the resulting genotype and phenotype via sequencing and drug resistance assays. This genetic transformation system offers a foundational tool for molecular studies of *Bd*, advancing our capacity to dissect molecular mechanisms of chytrid pathogenesis.

## INTRODUCTION

Amphibian populations are declining around the world, driven, in part, by the “frog-killing” chytrid fungus *Batrachochytrium dendrobatidis* (*Bd*) (1–4). Like other chytrid fungi, *Bd* has a biphasic life cycle that alternates between a motile dispersal form called a “zoospore” and a sessile reproductive form called a “sporangium” (5). Current infection models suggest that amphibian skin is colonized by zoospores that then invade epithelial cells and develop into intracellular sporangia whose growth and spread can rapidly lead to the death of the host by inhibiting electrolyte transport through the skin (5, 6). Fungal genotype, environmental factors, and host traits, such as species identity, life history, and skin microbiome are all thought to heavily influence *Bd* pathogenicity, making the outcome of infection difficult to predict (7, 8). Despite the clear importance to global ecology, testing these and other ideas about *Bd* cell biology, development, and pathogenicity have been limited by the lack of genetic tools with which to study the underlying molecular mechanisms.

To test key hypotheses about *Bd* pathogenesis, we recently developed a system for transient transformation by electroporation (9). Although this approach allows for transgene expression for up to four generations, it is inherently limited. First, electroporation of plasmids can result in overexpression artifacts due to the introduction of multiple plasmid copies or the use of non-native regulatory elements. Second, the use of extrachromosomal plasmids necessitates cloning entire open reading frames, which can be challenging for large proteins. Third, most aspects of *Bd* infection require multi-generational studies that go beyond the retention time of plasmids in the transient transformation system.

To overcome these limitations, we have developed a homology-mediated stable transformation system for *Bd* that facilitates sequence-dependent integration of transgene expression cassettes into its chromosomes. We use this system to (1) generate fluorescent protein fusions expressed from native loci, (2) use these fluorescent strains to visualize *Bd* infection on living animals, (3) test a key hypothesis about *Bd* development, and (4) develop a *Bd* gene knockout. These tools open the door to molecular hypothesis testing of *Bd* development, cell biology, and pathogenesis.

## RESULTS

### Stable genetic transformation of *Bd* by homologous recombination

To determine if stable transformation can be mediated by homologous recombination, we chose the glycolysis enzyme glyceraldehyde-3-phosphate dehydrogenase (GAPDH) as an initial target locus because its modification in other fungal and animal species does not induce detrimental phenotypes (10). To confirm that GAPDH is expressed in our lab strain JEL423, we performed RNA sequencing on zoospores and found that it is expressed at 1.3 ± 1.2 transcripts per million **(Fig. S1A)**, which is approximately the 42nd percentile of all genes. We also conducted immunoblot analysis on both zoospores and sporangia using anti-GAPDH antibodies and observed bands consistent with the expected size of *Bd* GAPDH in both samples (**Fig. S1B**). These data confirm that GAPDH is expressed across *Bd* life stages, making it an appropriate initial target for homologous recombination.

Having settled on the *GAPDH* locus as our target, we next engineered a plasmid designed to facilitate identification of transformants using two expression cassettes. The first cassette was designed to be appended to the 3’ end of the native *GAPDH* open reading frame and comprises the red fluorescent protein mRuby3 followed by the terminator region of the *Saccharomyces cerevisiae* alcohol dehydrogenase (*ScADH1ter*). The second cassette encodes the hygromycin B resistance marker hph under the control of the *Spizellomyces punctatus* histone 2B promoter (*SpH2Bpro*). The mRuby3 and the hygromycin resistance gene, along with the *ScADH1ter and SpH2Bpro* regulatory sequences, have been shown to be functional in *Bd* (9). To direct insertion of both cassettes at the 3’ end of the *GAPDH* gene, we flanked them with 1.35 kb-long homology arms with 100% sequence identity to the *GAPDH* locus. Successful homology-mediated chromosomal integration would therefore result in the expression of a GAPDH-mRuby3 fusion protein from the native *GAPDH* promoter as well as the hph protein from the heterologous *SpH2B* promoter (**Fig. 1A**).

**Figure 1:**
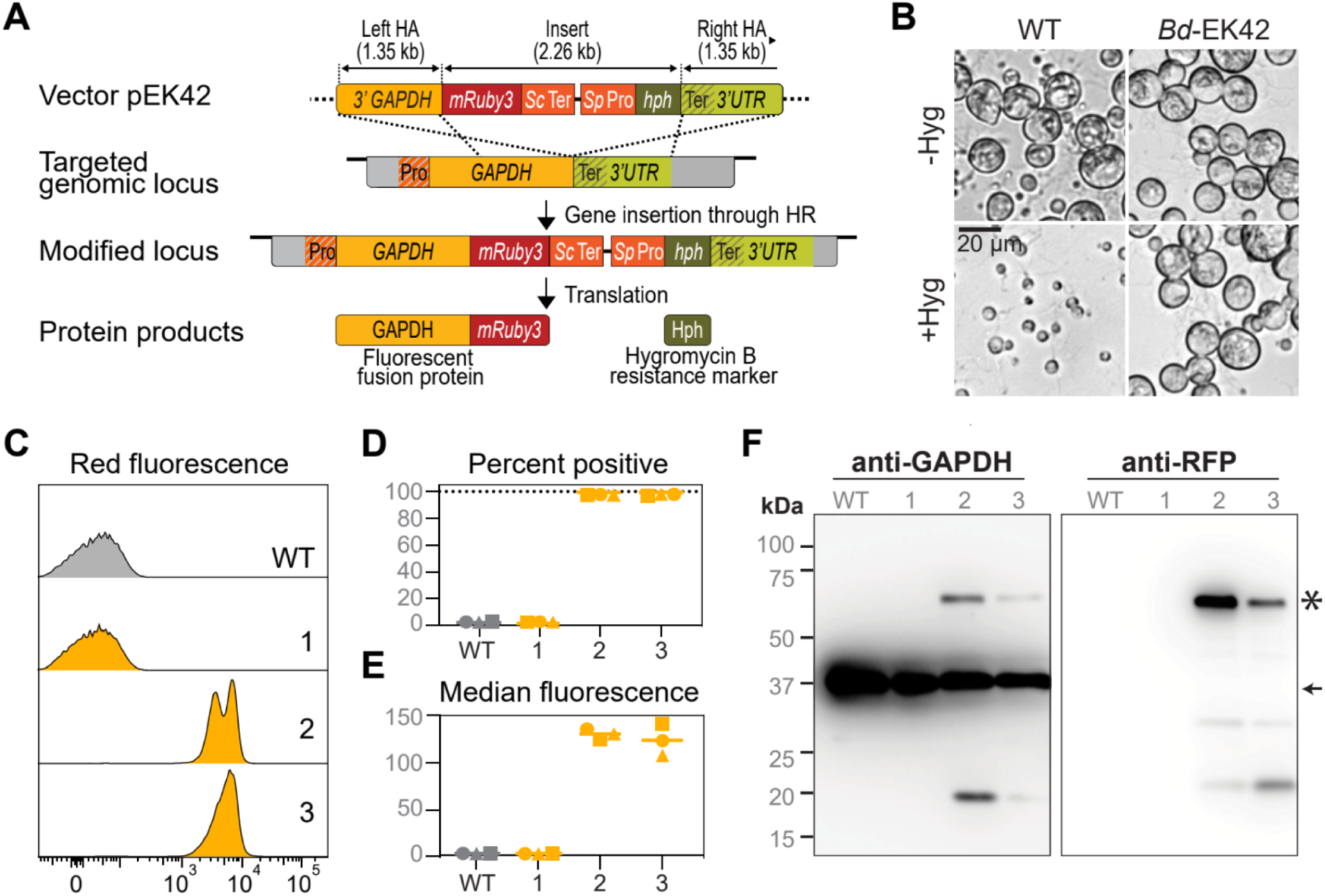
Generation of stable fluorescent *Bd* transformants through electroporation of a GAPDH targeting vector. (**A**) Schematic representation of the experimental strategy used to generate stable *Bd* transformants through homologous recombination. A flexible linker (GGGGS) was incorporated between the fluorescent protein and the protein of interest to prevent steric hindrance. To prevent off-target recombination events, we utilized non-endogenous regulatory sequences to confer/control expression of the transgenes (*Sc* terminator: *Saccharomyces cerevisiae ADH1* terminator, *Sp* promoter: *Spizellomyces punctatus H2B* promoter). *hph:* hygromycin resistance gene, HA: homology arm. (**B**) Bright field images of wild type cells (WT) and cells transformed with vector pEK42 (EK42-3) grown with or without hygromycin B for 3 days. (**C-E**) Measurement of mRuby3 expression in three single colony isolates (Bd-EK42-1,2,3) by flow cytometry. (**C**) Fluorescence distribution relative to the untransformed wild type strain (WT). Data presented is from one of three independent experimental replicates. (**D**) Percentage of mRuby3-positive cells from all three replicates. Replicates are represented by different symbols, and bars indicate means. Gates for mRuby3-positive cells were set using the wild type control for each replicate. (**E**) Median red fluorescence intensity of single cell events, with values normalized to the median intensities of wild type cells, which are set to 1 for each replicate. (**F**) Immunoblot analysis of GAPDH-mRuby3 expression in wild type cells (WT) and transformed cells (*Bd*-EK42-1,2,3) with anti-GAPDH (left) and anti-mRuby3 (right) antibodies. Star denotes expected band size for GAPDH-mRuby3 fusion protein (63 kDa) and arrow denotes expected size band size for native GAPDH protein (36 kDa).

We introduced the resulting circular plasmid (pEK42) into diploid *Bd* zoospores using our previously established electroporation protocol (9) and recovered the cells in non-selective liquid medium until they released daughter zoospores (see **Table S1** for an overview of the transformation protocol). We then transferred the zoospores onto agar plates containing 0.5 µg/ml hygromycin B to select for transformants. After 12-16 days of plate selection, we observed a few dozen active zoospores on the plate that continued to grow, resulting in a full lawn of sporangia after 20 days. We isolated single colonies from these lawns and successfully passaged the resulting cultures on selective agar every week for over four months (approximately 25 generations; **Fig. 1B**).

To determine if these hygromycin-resistant cells were also expressing mRuby3, we used flow cytometry to measure fluorescence in zoospores from three single-colony isolates (*Bd*-EK42-1, *Bd*-EK42-2, and *Bd*-EK42-3) (**Fig. 1C-E**). While wild type cells were nonfluorescent, nearly all cells in two of the three hygromycin-resistant strains were fluorescent, with median intensities 108-to 141-fold higher than wild type (**Fig. 1D-E**). For these two strains, two of the three biological replicates of the cytometry experiments showed bimodal levels of fluorescence. Using backgating, we found that the two peaks emerged from differently sized subpopulations as determined by forward scatter, likely representing motile and encysted zoospores (**Fig. S1D)**. The third strain (*Bd*-EK42-1), although hygromycin-resistant, contained almost no fluorescent cells (**Fig. 1D)**, with a fluorescence distribution indistinguishable from the wild type (**Fig. 1C**). To confirm that the fluorescence of the two positive strains is due to expression of GAPDH-mRuby3, we conducted immunoblot analysis of all three hygromycin-resistant single-colony isolates and the parental wild type strain using antibodies to both GAPDH and mRuby3 (**Fig. 1F**). The anti-GAPDH blot showed the expected band sizes of native GAPDH for all four strains and of the GAPDH-mRuby3 fusion for the two fluorescent strains. Similarly, the anti-mRuby3 blot confirmed bands of the expected size for the GAPDH-mRuby3 fusion for only the two fluorescent strains. Together, these data indicate that only two of the three hygromycin-resistant strains expressed the GAPDH-mRuby3 fusion.

To verify that the hygromycin-resistant transformants were generated by integration of the transformation vector, we analyzed the target site of wild type JEL423 and the three hygromycin-resistant strains using an integration-specific PCR screening strategy (**Fig. 2A**). The insert-specific primer pairs amplified fragments with sizes consistent with single-copy transgene insertions in the *GAPDH* locus of each transformant, as well as higher molecular weight fragments. No clear amplicons were detected in the wild type and plasmid controls, suggesting electroporation with the targeting vector resulted in transgene insertion in all three hygromycin-resistant lines.

**Figure 2:**
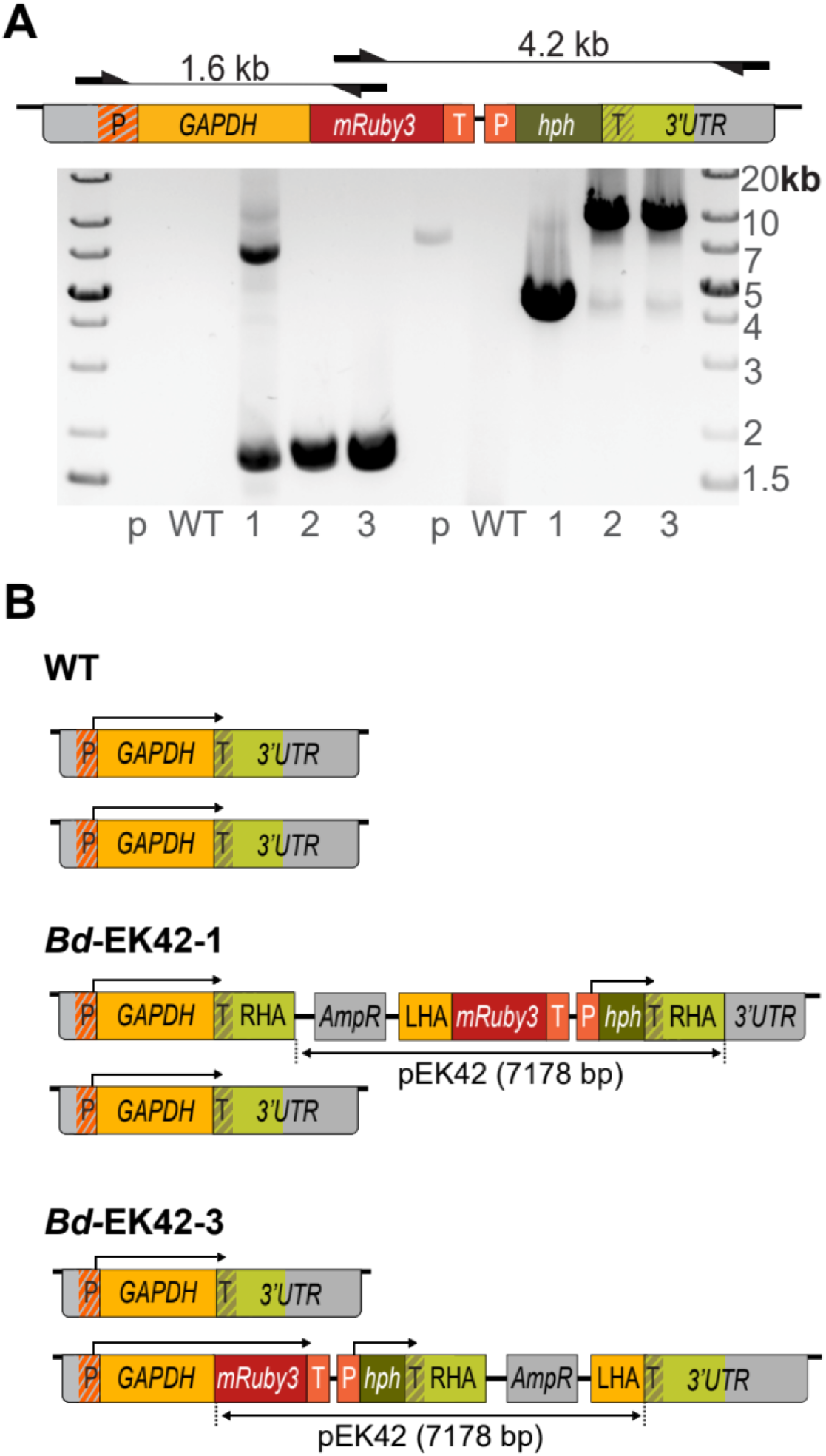
Stable transformants were generated by chromosomal integration of the introduced transformation vector into one allele. (**A**) (Top) Primer locations and expected amplicon sizes of different primer pairs specific to the transformed locus. Primers do not amplify wild type locus or plasmid alone. (Bottom) Gel electrophoresis of genotyping PCR confirming locus specific integration of expression cassettes into transformed clones *Bd*-EK42-1, 2 and 3 (WT=wild type, p=pEK42 plasmid control). (**B**) Diagram of reconstructed GAPDH alleles of WT and *Bd*-EK42-1 and 3 lines based on PCR and whole genome sequencing results. The arrows above each allele denote predicted open reading frames (P: promoter, T: terminator, *hph*: hygromycin resistance gene, HA: homology arm).

To further characterize the target loci in transformed *Bd* strains, we performed long-read whole genome sequencing of the parental strain (WT JEL423), the non-fluorescent transformant (*Bd*-EK42-1), and one of the two fluorescent transformants (*Bd*-EK42-3). We assembled haplotype-phased genomes for each strain and used SNPs and a 3-bp deletion to resolve two haplotypes at the wild type *GAPDH* locus (**Fig. S2**). Reads containing transgene sequences from the transformed clones showed identity to only one of the two haplotypes in each locus, consistent with insertions into a single *GAPDH* allele (**Fig. 2B, Dataset S3**). The fluorescent clone *Bd*-EK42-3 incorporated the entire targeting plasmid at the desired integration site—directly before the stop codon of the native *GAPDH* locus. In contrast, the non-fluorescent clone *Bd*-EK42-1 incorporated the plasmid immediately *after* the *GAPDH* stop codon, consistent with the lack of fluorescence in this strain. In lanes with multiple products, sequencing the high molecular weight bands previously detected by PCR (**Fig. 2A**) confirmed the accuracy of the inferred architecture of these loci. The lower molecular weight bands, however, were inconsistent with the whole genome sequencing results and are likely a PCR byproduct. The orientation of the plasmid-derived sequences relative to the native locus indicates that both transformants were generated by a single copy integration of the entire plasmid using only one homology arm, with the fluorescent strain using the left homology arm, and the non-fluorescent strain using the right homology arm. This type of whole plasmid integration by a single region of homology has been previously described in other model systems, including *E. coli* (11, 12) and yeast (13).

Taken together, these data confirm that our homologous recombination transformation system can stably introduce transgenic constructs to specific sites in the *Bd* genome. This approach can be used to fuse transgenic elements with endogenous proteins expressed from their native promoters. Moreover, because all three strains analyzed were clonally isolated from the same selection plate, single electroporations appear to result in multiple, independent, transformation events.

### Visualization of transgenic *Bd* in living frogs

Without methods to visualize infection on living animals, we cannot test key hypotheses about the spread of *Bd* in amphibian skin. To explore whether the GAPDH-mRuby3 fusion protein would be useful for visualizing *Bd* on live animals, we first imaged the fluorescent strain *Bd*-EK42-3 *in vitro* using fluorescence microscopy and observed cytosolic fluorescence at every growth stage (**Fig. 3A**), as well as punctate structures in zoospores (0 hours) and young sporangia (6 hours). This localization pattern is similar to the localization of GAPDH in other fungi, where it has been found in both the cytosol and intracellular compartments (14).

**Figure 3:**
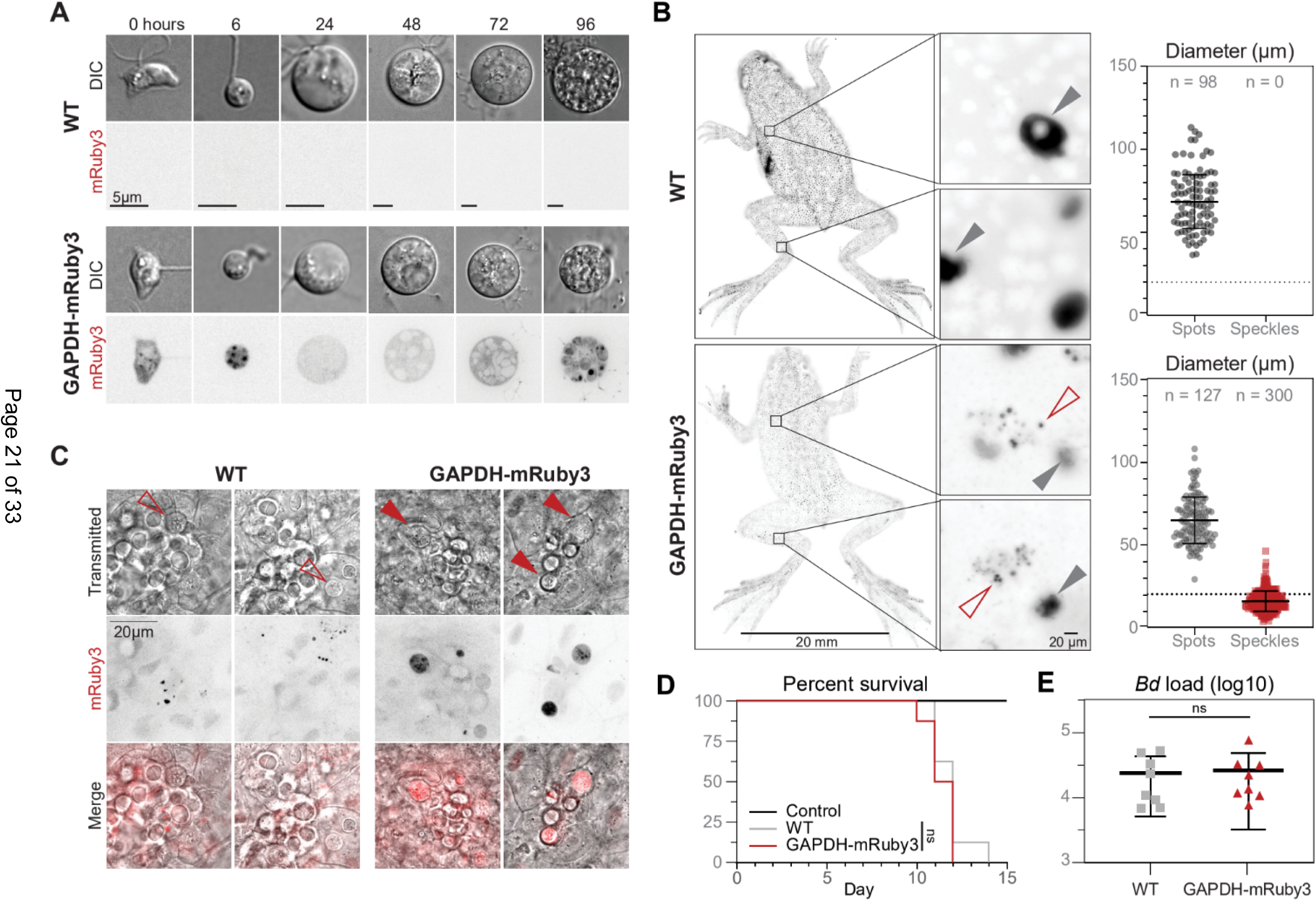
Cells expressing GAPDH-mRuby3 facilitate visualization of *Bd* infection on live frogs. **(A)** Representative single slice of spinning disc fluorescence microscopy showing GAPDH-mRuby3 localization throughout *Bd* life stages. **(B)** Widefield fluorescence microscopy of live adult *Hymenochirus boettgeri* frogs infected with either WT JEL423 or GAPDH-mRuby3 expressing cells. Fluorescent spots (gray arrowheads) are present across frogs infected with both WT and GAPDH-mRuby3. These spots are likely autofluorescent glands. Fluorescent speckles (empty red arrowhead) are present only in GAPDH-mRuby3 infected frogs. **(C)** Representative confocal images of webbing explants from adult *H. boettgeri* frogs infected with either WT or GAPDH-mRuby3. Sporangia can be observed growing in the epithelia in explants from both WT and GAPDH-mRuby3 infected frogs (empty and filled arrowheads). However, *Bd*-associated fluorescence is only observed in explants from GAPDH-mRuby3 infected frogs (filled arrowheads). **(D)** Survival curve of adult *H. boettgeri* frogs challenged with no *Bd* (control, n = 8 frogs), or 5 x 10^5^ zoospores of either the WT *Bd* strain JEL423 (n = 8 frogs) or the GAPDH-mRuby3 expressing strain (n = 8 frogs); WT:GAPDH-mRuby3 p=0.343. **(E)** Log-transformed *Bd* load per frog in each treatment at the conclusion of the experiment (when all treatment frogs died and control frogs were sacrificed). Frogs were swabbed and *Bd* DNA was probed for using qPCR and fitted to a standard curve of qPCR values from known zoospore amounts to establish pathogen load on the host; p=0.856.

Having confirmed that we could visualize GAPDH-mRuby3 using fluorescence microscopy, we next imaged the strain on living animals. We infected three adult African dwarf frogs (*Hymenochirus boettgeri*) with wild type JEL423 and another three African dwarf frogs with the GAPDH-mRuby3 strain. Imaging the dorsal and ventral surfaces of whole animals revealed large (66.3 ± 15.6 µm diameter) autofluorescent spots on both control and GAPDH-mRuby3 infected animals (**Fig. 3B**). In addition to these large spots, the GAPDH-mRuby3 infected animals also displayed speckles of 15.7 ± 6.0 µm diameter, consistent with the diameter of sporangia grown *in vitro* (**Fig. 3A**). The GAPDH-mRuby3 speckles were distributed across the body with no clear enrichment in any specific area (**Fig. 3B**). To obtain a more detailed understanding of how *Bd* interacts with amphibian skin cells, we turned to high resolution microscopy of webbing explants from the hind feet of these animals. While imaging with white light revealed large sporangia in all explants, only webbing from individuals infected with GAPDH-mRuby3 expressing cells showed *Bd*-specific fluorescence, with some images showing clear subcellular, punctate localization of the GAPDH-mRuby3 signal (**Fig. 3C**). These data indicate that we can use our transformation system to visualize subcellular structures in *Bd* growing within amphibian skin.

To explore whether altering the native *GAPDH* locus impacts *Bd* virulence, we conducted a challenge experiment in adult frogs. We inoculated adult *Hymenochirus boettgeri* with either wild type *Bd* (JEL423, n = 8), GAPDH-mRuby3 expressing *Bd* (*Bd*-EK42-3, n = 8), or no *Bd* as a control group (n = 8), and assessed host survival probability and *Bd* pathogen load. Inoculation with either wild type or GAPDH-mRuby3 expressing *Bd* resulted in host mortality within an average of 11 days (**Fig. 3D**), with no significant difference in virulence between the strains (log-rank test, χ² = 0.9, df = 1, p = 0.3). All animals inoculated with *Bd* showed signs of disease and died over the course of the experiment while no individuals in the control group showed signs of disease and tested negative for *Bd* at the end of the experiment (**Fig. 3E**). Moreover, the average *Bd* loads at the time of death did not differ between the two genotypes (one way ANOVA, F(_1,14_) = 0.078, *p* = 0.78). Taken together, these results show that transformed *Bd* strains expressing fluorescent proteins retain their virulence and can be detected on amphibians at least seven days post infection, allowing us to visualize progression of *Bd* infection on live animals and in skin explants.

### Use of endogenous tagging to test molecular hypotheses about chytrid development and pathogenesis

Having confirmed that we could use homologous recombination to precisely integrate exogenous DNA into *Bd* chromosomes, we next used this system to test molecular hypotheses regarding chytrid development. We focused on the assembly of chitin-containing cell walls, which are integral to the life cycle and pathogenicity of other fungi. Although the molecular mechanisms controlling cell wall synthesis in *Bd* are currently unknown, cell wall growth in yeast and filamentous fungi relies on delivery of vesicles carrying cell wall synthases to sites of cell wall assembly (15, 16). We previously hypothesized that a key event during chytrid development would be the release of chitin synthases from intracellular vesicles to the plasma membrane just prior to the *de novo* cell wall assembly that characterizes the zoospore-to-sporangia transition (17). We tested this hypothesis by determining the localization of chitin synthases during chytrid development, concentrating on a class of synthases known either as Myosin 17s or class IV chitin synthases. These chimeric proteins contain a myosin motor domain along with a chitin synthase 2 domain and have been implicated in the virulence of other fungal pathogens, including *Fusarium*, *Magnaporthe*, and *Ustilago* (18–21).

Myosin 17s are large, 200+ kDa proteins that are encoded by genes too long for previous plasmid-based expression methods. We therefore used our integration system to express a fluorescent Myosin 17. The JEL423 genome (GenBank GCA_000149865.1) encodes five Myosin 17s of which Myo17D has the highest transcript abundance in zoospores (**Fig. S1C**). Accordingly, we designed a plasmid to facilitate homology-mediated integration of mRuby3 at the 3’ end of the native Myo17D open reading frame, followed by a hygromycin selection cassette (see methods, **Table S1**). We transformed this plasmid into JEL423 and picked single-colony isolates. From these, we isolated two clones for subsequent analysis. Flow cytometry confirmed fluorescence in both single-colony isolates (**Fig. 4A-B**), with population-wide median fluorescence intensities 5-to 6-fold higher than those of wild type cells (**Fig. 4C**). We confirmed genomic integration using insert-specific PCR, observing bands consistent with single-copy transgene insertions at the *Myo17D* locus of each transformant (**Fig. 4D**). Unlike for the *GAPDH* insertion, we did not detect additional higher molecular weight bands, hinting that the expression cassette but not the plasmid backbone was inserted. Whole genome sequencing of the isolate with the greatest fluorescent expression, *Bd*-EK48-2, confirmed the intended integration in a single Myo17D allele (**Fig. 4E, Dataset S3**).

**Figure 4:**
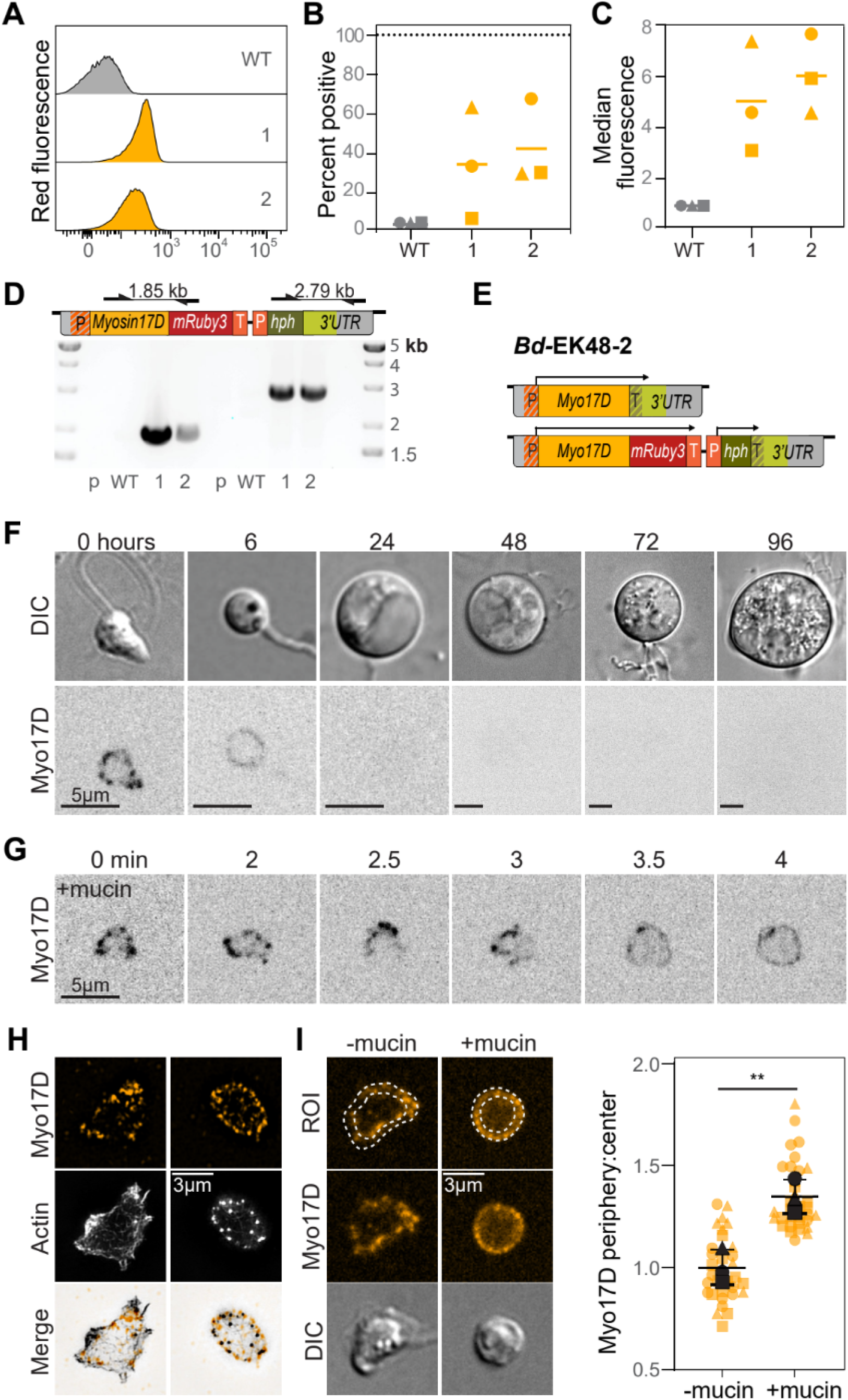
Stable Myo17D-mRuby3 expression provides novel insights into dynamics/molecular mechanisms of *Bd* development. (A-C) Measurement of mRuby3 expression in three single colony isolates (*Bd*-EK48-1,2,3) by flow cytometry. Wild type data is the same as shown in Figure 1C-E. **(A)** Fluorescence distribution measured by flow cytometry, relative to untransformed wild type strain. Data shown is from one of three independent experimental replicates. **(B)** Percentage of mRuby3-positive cells from all three replicates. Replicates are represented by different symbols; bars indicate means. Gates for mRuby3-positive cells were set using the wild type control for each replicate. **(C)** Median red fluorescence intensity of single cell events, with values normalized to the median intensities of wild type cells, which are set to 1 for each replicate. **(D)** (Top) Primer locations and amplicon sizes of different primer pairs specific to the transformed locus. Primers do not amplify the wild type locus or plasmid alone. (Bottom) Gel electrophoresis of genotyping PCR confirming locus specific integration of expression cassettes into transformed clones *Bd*-EK48-1 and 2 (WT=wild type, p=pEK48 plasmid control). **(E)** Diagram of reconstructed Myo17D alleles of *Bd*-EK48-2 based on PCR and whole genome sequencing results. The arrows above each allele denote predicted open reading frames. **(F)** Representative single slice of spinning disc fluorescence microscopy showing Myo17D-mRuby3 localization throughout *Bd* life stages. **(G)** Zoospore expressing Myo17D-mRuby3 undergoing mucin-induced encystation. **(H)** Structured illumination microscopy of *Bd* zoospore (left) and encysted zoospore (right) expressing Myo17D-mRuby3 (magenta), fixed and stained for actin with Alexa Fluor 488 Phalloidin (green). **(I)** (Left) Representative cells showing cells traced using phalloidin staining as a guide and divided into two concentric equal areas: the cell periphery and cell center. (Right) Ratio of Myo17D-mRuby3 fluorescence in the cell periphery to the cell center. Fluorescence was normalized to background fluorescence; unpaired t-test **p=0072.

To determine whether and how the subcellular distribution of Myo17D changes during chytrid development, we examined its localization at each life stage. We found that Myo17D-mRuby3 is primarily expressed in early stages of *Bd* development, forming punctate structures in zoospores, while displaying a more uniform distribution along the cell periphery of young sporangia (**Fig. 4F**). To explore the dynamics of this shift in localization, we treated zoospores with mucin to induce encystation—the transition from non-walled zoospores to cell wall encased sporangia (17). In cells undergoing encystation, Myo17D-mRuby3 transitioned within minutes from its punctate to its peripheral distribution (**Fig. 4G**), a sequence consistent with a rapid movement of Myo17D from intracellular vesicles to the plasma membrane. To explore this idea further, we stained Myo17D-mRuby3 expressing cells for actin, which localizes to the periphery in both zoospores and sporangia. In zoospores, actin forms a layer under the plasma membrane, while in sporangia it assembles into bright puncta at the cell edge (22).

Super-resolution imaging of stained zoospores revealed the Myo17D-mRuby3 puncta localized within the volume enclosed by the peripheral actin shell. In contrast, Myo17D-mRuby3 in early sporangia was interspersed with actin puncta found at the cell edge (**Fig. 4H**). To quantify this observation, we calculated the ratio of peripheral to internal Myo17D-mRuby3 fluorescence and confirmed an increase of signal ratio from 1.1 ± 0.1 in un-walled zoospores to 1.4 ± 0.1 in early sporangia (**Fig. 4I**). Taken together, these data indicate that chytrid development involves a rapid deployment of chitin synthase from interior puncta to the cell periphery during *de novo* cell wall assembly, consistent with our previously proposed molecular hypothesis (17).

### Homology-mediated knockout of the *Bd URA3* gene

Finally, we extended our use of homologous recombination to generate targeted knockout lines of the *URA3* locus, which encodes the enzyme orotidine-5′-phosphate decarboxylase (ODCase), a key component of the uracil synthesis pathway (23). Deletion of this gene in yeast enables negative selection using the uracil analog 5-fluoroorotic acid (5-FOA) that is converted into a toxin in the presence of ODCase (23).

Our initial attempt at generating a *URA3* knockout employed a two-step approach intended to sequentially replace both copies of the *URA3* coding sequence. We first electroporated wild type cells with a construct designed to replace *URA3* with hygromycin resistance and isolated hygromycin-resistant colonies. We then electroporated one of these colonies with a second *URA3*-targeting construct with the same homology arms but encoding blasticidin resistance, and isolated double-resistant colonies (**Fig. S3A**). To determine whether these lines had functional ODCase activity, we grew them in the presence of 5-FOA and confirmed that, while no WT sporangia and very few hygromycin-resistant sporangia (0.76% ± 0.01%) showed robust growth in 5-FOA, the double-resistant line grew in 5-FOA, indicating disruption of ODCase activity (**Fig. S3B**). To determine the genotype of this 5-FOA resistant line, we conducted insert-specific PCR and whole genome sequencing. We recovered only one allele of the *URA3* locus, which contained multiple, complete tandem insertions of both targeting plasmids (**Fig. S3C, S3D, Dataset S3**). Although this first approach was successful at generating a *URA3* knockout line, the resulting genome architecture suggests it was a product of homologous recombination combined with a loss of the heterozygosity event, and not a product of sequential replacement as intended.

Because our first approach resulted in multiple insertions, we hypothesized that *Bd* could undergo multiple rounds of homologous recombination from a single electroporation event. Therefore, we next attempted to generate a *URA3* knockout line using a one-step approach. This time, we linearized the targeting plasmid encoding hygromycin-resistance prior to electroporation to prevent integration of the plasmid backbone (**Fig. 5A**). We electroporated wild type cells with this linearized construct, then isolated hygromycin-resistant colonies. We screened these colonies by insert-specific PCR and confirmed successful integration of the hygromycin resistance gene into the *URA3* locus. We also amplified the *URA3* coding sequence from a subset of the hygromycin-resistant colonies, indicating these lines were likely heterozygous knockout lines (**Fig. 5B**). Whole genome sequencing of one of the other lines from which we could not PCR amplify the *URA3* coding sequence recovered two distinct alleles of the *URA3* locus, with the coding sequences of both having been replaced with the hygromycin resistance gene as intended (**Fig. 5C, Dataset S3**).

**Figure 5:**
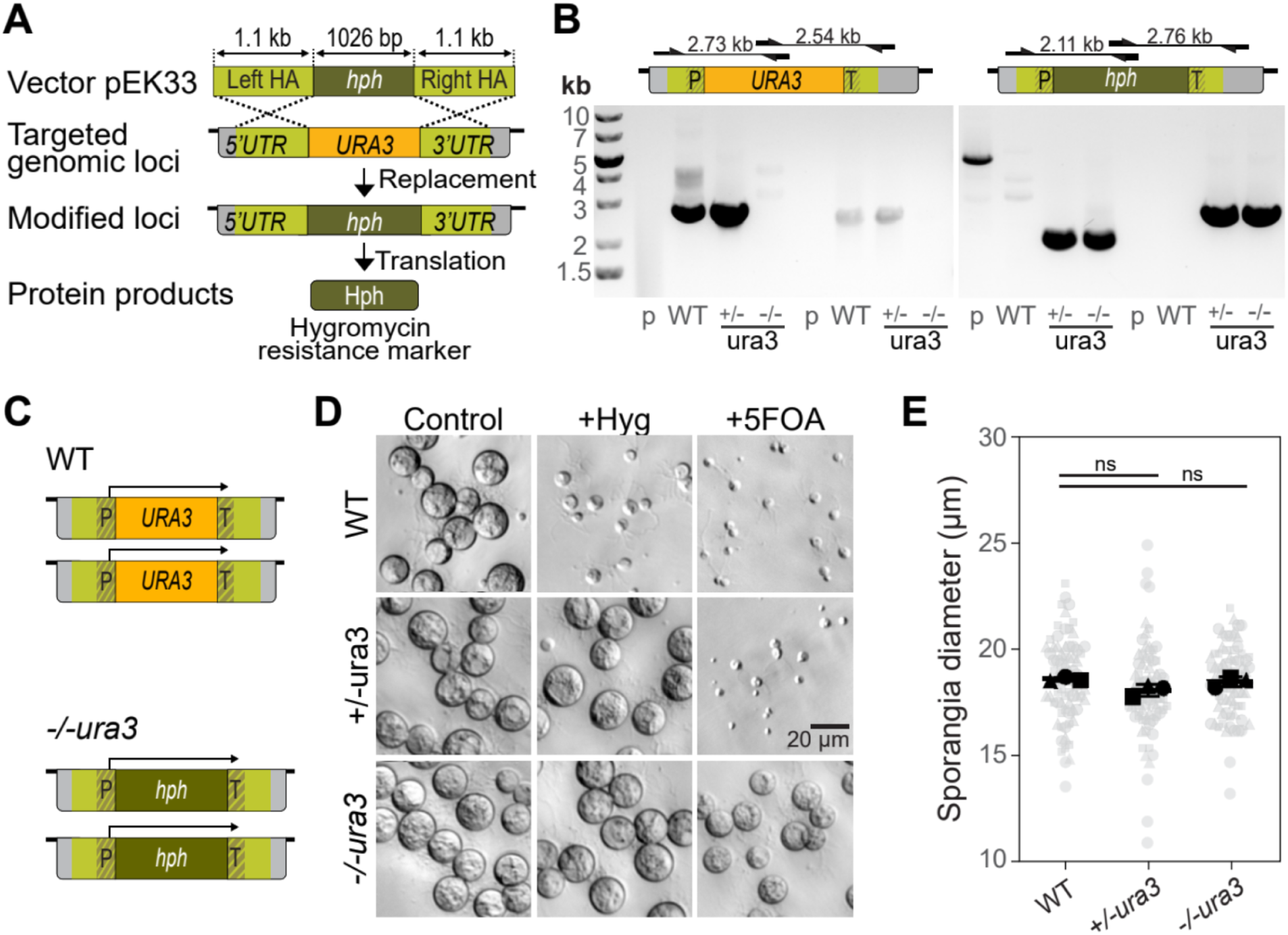
Gene replacement by homologous recombination enables the generation of knockout strains. **(A)** Schematic representation of the experimental strategy used to generate stable *URA*3 knock-out through gene replacement using homologous recombination. **(B)** (Top) Diagrams showing primer locations and amplicon sizes of different primer pairs to native *URA3* locus (left) or *hph* replaced locus (right). (Bottom) Gel electrophoresis of genotyping PCR confirming presence of the native *URA3* locus (left) and *hph* locus (right) specific integration into WT, +/-*ura3*, and -/-*ura3* lines (p = pEK33 plasmid control). **(C)** Diagram of reconstructed *URA3* alleles of WT and *-/-ura3 cell* lines based on whole genome sequencing assembly results (hph: hygromycin resistance gene, HA: homology arm, P: promoter, T: terminator). **(D)** Bright field images of WT, +/-*ura3*, and -/-*ura3* lines cells grown for 3 days in tryptone media supplemented with hygromycin B or 5-FOA. **(E)** Sporangia diameter of WT, +/-*ura3*, and -/-*ura3* cells grown in tryptone media for 3 days.Unpaired t-test WT:+/-*ura3* p=0.071; WT:-/-*ura3* p=0.486.

Having confirmed the genotype of the homozygous knockout, we next assayed its phenotype. While the heterozygous *URA3* deletion showed no growth in 5-FOA, the homozygous *URA3* knockout was resistant, indicating that we had successfully disrupted ODCase enzymatic activity (**Fig. 5C**). Finally, because disruption of the *URA3* gene in other species can result in developmental defects (24), we quantified the growth of *Bd URA3* knockout lines. We grew cells in non-selective media and measured the diameters of sporangia after 3 days and found no significant difference between WT, heterozygous, and homozygous knockout cells (**Fig. 5E**). Taken together, these data indicate that our single-step approach can be used to generate a clean knockout line.

## DISCUSSION

Here we establish homology-mediated stable transformation of *Bd* to facilitate precise insertion of exogenous DNA into the *Bd* genome. We use this system to tag endogenous proteins, generate *Bd* knockout strains, and use these new tools to investigate molecular mechanisms driving chytrid pathogenesis and development.

Although chytridiomycosis is well established as a key driver of global amphibian decline, the mechanisms that *Bd* uses to infect its host are largely unknown. Here we show that we can use homology-mediated transformation to develop *Bd* strains with stable fluorescence that can be visualized on live animals. Fluorescent imaging of *Bd* infection offers a powerful tool to track and understand fungal invasion patterns, tissue colonization, and host response timing. These insights could contribute to developing targeted strategies to mitigate the impact of chytrid disease on amphibian populations. Moreover, because these strains stably express fluorescent protein, they can be readily shared with other labs who study *Bd* pathogenesis but who are not equipped for molecular biology. To promote such collaborative studies, we have deposited representative strains in the CZEUM repository, where they are now readily available to the research community (**Table S3**, https://czeum.herb.lsa.umich.edu/).

We also used homologous recombination to develop strains to test a previous hypothesis that *Bd* encystation involves rapid deployment of chitin synthases to the plasma membrane (17). Indeed, visualization of the tagged chitin synthase Myo17D showed rapid reorganization from intracellular puncta to the cell periphery upon encystation. Furthermore, Myo17D expression was primarily restricted to early sporangial development, raising the possibility that other Myosin 17 genes may be sequentially expressed or play distinct roles in cell wall assembly. Future work on other Myosin 17s could provide insight into the coordination of this multigene family during *Bd* development.

Finally, we have reached a major milestone by achieving targeted gene deletion in *Bd*. We generated a *URA3* knockout line and used whole genome sequencing to confirm the genetic identity of the mutant, as well as confirming its phenotype using drug resistance. This work lays the foundation for developing targeted gene knockouts to test key hypotheses about the role of specific genes in *Bd* pathogenesis, such as those encoding chitin-binding proteins (25), Crinkler effector proteins (26), and metalloproteases (27).

This transformation system could be enhanced by optimizing the design of targeting constructs. Although we have demonstrated recombination using long (1-1.5 kb) homology arms, homologous recombination in other fungi can use arms as short as 60-600 base pairs (28–30). Therefore, determining the required length for homology arms in *Bd* could be used to reduce the size of targeting constructs. Furthermore, while some clones integrated only the intended cassette, other clones integrated the entire plasmid. Not only did whole plasmid integration give unexpected genome architectures, screening these clones via PCR proved unreliable, as the reactions produced additional bands which were inconsistent with whole genome sequencing. Our success with using a linearized construct to generate a clean *URA3* knockout suggests that this approach may reduce the frequency of unintended integration products. Finally, because the current method for isolating single clones is time and labor intensive, developing new methods for colony isolation, such as by flow cytometry, could further streamline the transformation process, as well as allow for quantifying the rates of homologous recombination in *Bd*.

In summary, by harnessing endogenous homologous recombination pathways, we have created a system for homology-based integration that facilitates stable transformation, endogenous tagging, and gene deletion in *Bd*. These tools substantially advance functional genetics for the study of *Bd* biology, which can inform strategies to mitigate its impacts on global amphibian populations. They can also serve as a foundation for additional technologies,such as inducible expression systems, CRISPR-based gene modifications, and genetic screens. To promote the use of these tools by the broader research community, we have deposited all constructs at Addgene (**Dataset S1**) and submitted key strains to the CZEUM fungal collection. We look forward to seeing how the field utilizes these resources in future studies.

## MATERIALS AND METHODS

### Cell culture

We used the *Batrachochytrium dendrobatidis* strain JEL423 (from the highly virulent global panzootic lineage) for all experiments. To obtain cells adapted to growth on solid media, cryopreserved wild type *Bd* cultures were revived as described previously (31) and maintained in liquid 1% tryptone (w/v) media at 24°C in tissue culture-treated flasks (Fisher, Cat. No. 50-202-081) until they grew at a 3 day lifecycle. We then collected synchronized *Bd* zoospores in media, strained them through sterilized 25 mm Whatman Grade 1 filter paper (Cat. No. 1001–325) to remove sporangia and cell clumps, centrifuged them for 5 min at 2,500 × g at room temperature (∼22°C), and resuspended the cell pellet in Bonner’s salts (10.27 mM NaCl, 10.06 mM KCl, 2.7 mM CaCl_2_ in ultrapure water) to a concentration of ∼6 × 10^5^ cells/ml. Next, we spread 900 µl of the cell solution onto 1.5% agar plates containing 1% tryptone (Sigma, Cat. No. T7293), dried the plates with the lids open for 10 minutes, sealed them with Parafilm, and transferred them to an incubator set to 24°C. In our experience, cells cultured in this manner release the first generation of daughter zoospores after 4-5 days, followed by the second generation after 7-8 days. We continuously grew the cells, transferring them to new plates every 8 days for at least 6 generations (2 transfers), before considering them adapted to growth on solid media and suitable for transformation.

For regular maintenance of strains grown on agar plates, we induced the release of zoospores by flooding actively growing plates with 1.5 ml of Bonner’s salts, incubating them for 2 h at room temperature (∼22°C), then spreading 25-50 µl of this cell solution and 850 µl Bonner’s salts onto plates supplemented with antibiotics as required (0.5 µg/ml hygromycin B [Gibco, Cat. No. 10687010], 10 µg/ml blasticidin S [Fisher scientific, Cat. No. A1113903], 1% tryptone, 1.5% agar). We then dried, sealed, and incubated the plates at 24°C as described above for wild type cells.

### Vector construction

We used the Phusion high-fidelity DNA polymerase (New England Biolabs, Cat. No. M0530L) for all PCR reactions. To aid in the design of homology arms and identify suitable primer pairs for transformant screening, we verified target locus sequences by PCR amplification from JEL423 genomic DNA (**Dataset S2**) and subsequent long-read sequencing at plasmidsaurus (SNPsaurus LLC). Fungal regulatory sequences (*ScADH1ter*, *SpH2Bpro*) and *Homo sapiens* codon-optimized reporter genes (*mRuby3*, *hph*) were demonstrated to be functional in *Bd* (*9*). Gene targeting vectors were synthesized by either Twist Bioscience (South San Francisco, United States) or generated by Sequence and Ligation Independent Cloning (32, 33). We verified the sequence of all constructs by whole-plasmid sequencing (SNPsaurus LLC). For details about each plasmid used in this study, refer to **Dataset S1**. Complete plasmid sequences and plasmids have also been deposited at Addgene (https://www.addgene.org/Lillian_Fritz-Laylin/, **Dataset S1**).

For transformation experiments, we performed large-scale preparations of reporter plasmids with a Midiprep kit (NucleoBond Xtra Midi, Machery-Nagel, Cat. No. 740410.50) and resuspended the DNA pellet in low TE buffer (10 mM Tris, 0.1 mM EDTA, pH 8.0) to a concentration of at least 2.7 µg/µl.

To linearize plasmids pEK33 and pSB02 for transformation, we incubated 100 µg of plasmid DNA with 100 units of restriction enzyme SspI-HF (New England Biolabs, Cat. No. R3132S) at 37°C for 18 hours, then heat inactivated by incubation at 65°C for 20 minutes. To precipitate the DNA, we added 0.1 volumes of 3 M sodium acetate (pH 5.2) and 1 volume 100% ice-cold ethanol and stored at -80°C for 30 minutes, then centrifuged for 30 minutes at 20,000 × g at 4°C and washed the pellet twice with 70% ethanol by centrifugation for 5 minutes at 20,000 × g at room temperature. Finally, we resuspended the DNA pellet in low TE buffer and checked for complete plasmid digestion by gel electrophoresis.

### Electroporation

We used the electroporation protocol developed for transient transformation (9) with the following modifications: (**1**) To adapt *Bd* cells to growth on plates for downstream single-colony isolation protocols, cells were grown on and harvested from agar plates instead of from liquid cultures. To induce the release of zoospores, we flooded actively growing plates with 1.5 ml of Bonner’s salts and incubated them for 2 h at room temperature (∼22°C). We then collected the cell solution and removed sporangia and cell clumps by straining through a sterilized 25 mm Whatman Grade 1 filter paper (Sigma, Cat. No. 1001–325). A standard 94/16 mm petri plate of healthy, densely-grown *Bd* yields enough zoospores for 4 to 5 electroporation cuvettes. (**2**) Across independent transformation experiments, the working concentration of the zoospore suspension ranged from 1.5 × 10^7^ to 3.25 × 10^7^ cells/ml, resulting in the loading of 0.6 to 1.3 × 10^6^ cells into each electroporation cuvette. (**3**) We used 40 µg of circular or linearized expression vector in each electroporation. (**4**) To test for successful plasmid delivery, we allowed the cells to recover for 18 h at 24°C, and then subjected one 600 µl aliquot of electroporated cells (about 1/7 of the sample) to early selection pressure as described in (9). We then incubated the cells for an additional 78 h at 24 °C before assessing their survival by light microscopy. (**5**) For specified transformations, we first linearized the plasmid before electroporation to try to improve transformation efficiency and avoid integration of the plasmid backbone.

### Isolation of stable transformants

For selection of stable transformants, we grew the electroporated cells at 24°C until at least 70% of sporangia had released zoospores (approx. 96 h after electroporation), then pooled the media and cells from all wells of a sample into a single 50 ml centrifuge tube and pelleted the cells by centrifugation for 6 min at 2,500 × g at room temperature. We then carefully decanted the supernatant, added 900 µl of Bonner’s salts to the pellet and remaining liquid, and gently resuspended the cell pellet, spread 900 µl of the cell solution on antibiotic selection plates, mixed the remaining volume with an additional 750 µl of Bonner’s salts, and spread it on survival control plates (plain 1% tryptone solid media). We next dried the plates with the lids open for 12-15 minutes, sealed them with parafilm, and transferred them to a 24 °C incubator. To facilitate the growth of a healthy cell culture, it is crucial to ensure that the plates are constantly covered by a thin layer of liquid. To ensure this, we checked plates daily and added additional Bonner’s salts as needed to maintain the liquid layer. During the selection period, we inspected the plates every four days using a tissue culture microscope, assessing the number and mobility of zoospores. In some transformants, after 4-5 days of growth on selection plates, we observed a burst of zoospore release, potentially corresponding to a period of transient expression of the antibiotic resistance protein (9); after 10-12 days, growth slowed and we observed only a handful of motile zoospore–less than 100 swimming zoospores per field of view. After 3-6 weeks of incubation, if the transformation was successful, lawn growth with 100-1000 swimming zoospores per field of view is detectable. Transformation outcomes, including the appearance of antibiotic-resistant and fluorescent cells, varied among individual experiments and plasmids (**Table S2**).

To isolate clonal transformant strains, we performed colony isolation. To obtain single colonies of *Bd*, it is important to use freshly poured agar plates to avoid desiccation, break down sporangial clumps by pipetting, and let the liquid absorb enough to prevent liquid pools that facilitate zoospores swimming across the plate. We harvested cells by flooding the initial selection plate with Bonner’s salts and incubating for 2 h at room temperature. We then collected the cell solution, diluted it 1:250-1:500 in Bonner’s salts, and spread 800 µl of the dilution on a fresh selection plate. The plate was allowed to dry with the lid open until there was no visible liquid layer covering the surface (approximately 20 minutes on a freshly poured plate), then returned to the 24 °C incubator. Once colonies appeared, we picked them with a pipette tip and resuspendend each in 100 µl of Bonner’s salts, gently breaking them up by pipetting. Next, we added 2 µl of the cell solution and 800 µl of Bonner’s salts to a new antibiotic selection plate and repeated the colony isolation procedure. After the second round of single colony isolation, we transferred the entire colony suspensions onto selective plates to amplify each clonal culture for cryopreservation (31) and downstream analysis.

For an overview of a typical timeline for the entire transformation process, including advance preparation of required reagents and materials, see **Table S1**. Strains listed in **Table S3** were deposited in CZEUM and are also available from the Fritz-Laylin lab upon request.

### Antibiotic resistance assay

To assay antibiotic resistance of lines, 5 × 10^5^ cells in 1 ml 1% tryptone media supplemented with 0.5 µg/ml hygromycin B, 50 µg/ml blasticidin S, or 1 mg/ml 5-fluoroorotic acid (ThermoFisher Scientific, Cat. No. R0811) were seeded in wells of a 24-well tissue culture plate (Fisher Scientific, Cat. No. 50-202-139). We then sealed the plates with parafilm and incubated at 24°C for 3 days before imaging.

### Mucin-induced encystation

We resuspended 20 mg/ml mucin (Sigma, Cat. No. M1778) in Bonner’s salts then centrifuged at 15,000 × g for 10 minutes to remove particles. We then adhered zoospores resuspended in Bonner’s salts to a 96-well plate coated with 0.5 mg/ml Concanavalin A and treated with mucin at a final concentration of 10 mg/ml.

### Phalloidin staining

We adhered zoospores resuspended in Bonner’s salts to a 96-well plate coated with 0.5 mg/ml Concanavalin A for 5 minutes. We then treated cells with Bonner’s salts or mucin (Abcam, Cat. No.ab144291) for 5 minutes before fixing cells in 4% PFA, 50 mM sodium cacodylate, pH 7.2 on ice for 20 min, and washing them thrice with PEM (100 mM PIPES, pH 6.9, 1 mM EGTA, 0.1 mM MgSO_4_). We permeabilized and stained the cells with 66 nM Alexa Fluor 488 Phalloidin (ThermoFisher Scientific, Cat. No. A12379) in 0.1% Triton X-100 PEM buffer for 20 min at room temperature, and washed three times with PEM.

### Microscopy

Images in **Figs. 1B, 5D** and **S3B** were taken with a Nikon Ti2-E inverted microscope equipped with a Photometrics Prime BSI Express camera and 40x Plan Fluor 0.6 NA objective, controlled through NIS-Elements software (Nikon). Images were acquired using Phase-contrast with LED transmitted light. For experiments shown in **Figs. 3A**, **4F**, **4G**, and **4I**, we used a Nikon Ti2-E inverted microscope equipped with a Crest X-Light spinning disk, a Photometrics Prime 95B camera, and a Lumencor Celesta light source for fluorescence, controlled through NIS-Elements AR software (Nikon). Images were taken using a Plan Apo λ 100x 1.45 NA oil objective in DIC (acquired using transmitted light) and fluorescence microscopy with excitation light 546 nm laser to visualize mRuby3 and 488 nm laser to visualize Alexa Fluor 488 Phalloidin.

For images in **Figs. 3A and 4F**, the “0 hour” time point was taken by harvesting and filtering zoospores from plates as previously described, and resuspending cells to a working concentration of 10^5^ cells/ml in Bonner’s salts. We then adhered zoospores to a 96-well plate (MatriPlate, Cat. No. MGB096-1-2-LG-L) coated with 0.5 mg/ml Concanavalin A (Sigma, Cat. No. C2010) for 5 minutes. To image time points at 6 hours and beyond, we resuspended harvested zoospores to a working concentration of 5 × 10^4^ cells/ml in 1% tryptone media. We then seeded 2 ml of resuspended cells into each well of a 12-well glass-like plate (Cellvis, Cat. No. P121.5P) and returned the plate to a 24 °C incubator. Before imaging, we washed wells three times with Bonner’s salts.

Super-resolution structured illumination microscopy (SIM) imaging in **Fig. 4H** was performed using Nikon TiE microscope equipped with a Hamamatsu Photonics C11440-22C sCMOS camera and SR Apo TIRF 100X 1.49 NA objective, controlled through NIS-Elements AR software (Nikon). Fifteen raw images were acquired with five phases of parallel grid pattern for each of three grid rotations to reconstruct one SIM image. Quality of raw images was evaluated using SIMcheck (34). Images were analyzed and reconstructed using NIS-Elements AR.

### Flow cytometry

To measure fluorescence of zoospores, we harvested wild type and transformant cells grown for 5 to 6-days on plates, then filtered them through a sterilized 25 mm Whatman Grade 1 filter paper to remove sporangia. We then washed and fixed the cells using the protocol previously used for transient fluorescent transformants (9). We conducted all flow cytometry measurements on a BD Dual LSRFortessa cell analyzer (BD Biosciences), using the 561 nm laser to detect mRuby3 fluorescence and recording a minimum of 13,900 single cell events per sample. We analyzed the resulting data using FlowJo v10.9, setting gates based on events recorded from wild type cells.

### DNA isolation

To isolate high-molecular-weight (average DNA fragment size ≥ 20 kb) *Bd* genomic DNA, we used a fungal cetyltrimethylammonium bromide (CTAB)/chloroform extraction protocol. To obtain ∼15 µg gDNA, we flooded two confluent plates of a single *Bd* strain with 2 ml of Bonner’s salts and incubated them at room temperature for 2 h. We then dislodged adherent sporangia with a cell scraper (Fisher Scientific, Cat. No. 50-201-974) and collected all cell material (zoospores and sporangia) into a single 50 ml centrifuge tube, raising the volume to 20 ml with Bonner’s salts. Following this, we washed the cells once with 10 ml of Bonner’s salts by centrifugation at 3,000 × g for 5 min at room temperature, resuspended them in 1.5 ml of fresh Bonner’s salts, transferred them to a fresh 2 ml microcentrifuge tube, centrifuged them again at at 3,000 × g for 5 min at room temperature, and removed the supernatant. We then resuspended the pellet in 900 µl of freshly-made CTAB/PVP buffer (100 mM Tris pH 8.0, 2 M NaCl, 10 mM EDTA pH 8.0, 1% CTAB; 1% PVP; 1% β-Mercaptoethanol) prewarmed to 65°C and transferred the sample to a new 2 ml centrifuge tube. Next, we vortexed the tube for 30 s, followed by a 30 min incubation at 65 °C and another 30 min incubation at 24°C, both at 400 rpm in a Thermomixer R (Eppendorf). After a 5 min incubation on ice, we removed proteins, lipids, and other cellular debris from the DNA thrice with an equal volume of chloroform (Fisher Scientific, Cat. No. C298-4), followed by centrifugation at 20,000 × g for 10 minutes at 4°C. To remove contaminant RNA, we then collected the aqueous phase and treated it with 400 µg RNase A (ThermoFisher Scientific,Cat. No. EN0531) for 30 min at 37°C. To precipitate the DNA, we subsequently adjusted the salt concentration with 0.1 volumes of 3 M sodium acetate (pH 5.2), added 2 µl glycogen (Invitrogen, Cat. No. 10814010) and 0.7 volumes isopropanol, mixed by tapping the tube, and centrifuged the sample immediately at 20,000 × g for 35 min at 4°C. We carefully removed the supernatant with a pipette, washed the pellet twice with 1.5 ml of 70% ethanol by centrifugation at 20,000 × g for 10 min at room temperature, and resuspended the air-dried pellet in 70 µl of low TE buffer. Following overnight incubation at 4°C, we determined the DNA integrity and quantity by gel electrophoresis and spectrophotometry (NanoDrop 2000, ThermoFisher Scientific).

### PCR genotyping

Isolated clones were screened for transgene integration into target loci by PCR using primers pairs specific to transgene insertion (**Dataset S2**). Primers were designed to amplify sequences upstream and downstream of vector homology arm sequences. Transformed clones were identified by expected amplicon size after DNA gel electrophoresis.

### Genome sequencing and assembly

Whole genome sequencing of the -/-*ura3* knockout line was performed by Plasmidsaurus with Oxford Nanopore Technology PromethION R10.4.1. We obtained 3.8 billion total bases sequenced with ∼160x coverage.

For all other sequenced lines, whole genome sequencing was performed by Azenta Life Sciences (South Plainfield, NJ, USA). DNA quantity and quality were assessed using a Qubit fluorometer (Invitrogen) and TapeStation system (Agilent). DNA was sheared using a Megaruptor 3 system, and PacBio SMRTbell libraries were prepared following the manufacturer’s protocol. Sequencing was performed on a PacBio Revio platform using v3.0 chemistry in circular consensus sequencing (CCS/HiFi) mode for 24 hours. We obtained at least 5.7 billion CCS bases per sample (∼230x coverage), with mean read lengths of 6.6–11.4 kb, mean quality scores ≥ 0.999, and 14–22 passes per read.

Genome assembly and analysis were performed using workflow and tools described in (35). Briefly, de novo genome assembly was performed using Hifiasm v0.19.8 (36) with default parameters, generating haplotype-phased assemblies. Assembly quality was assessed using QUAST v5.0.0 (37) with default parameters, reporting total assembly size (range: 26–34 Mb), number of contigs (75–403), N50 (1.2–1.8 Mb), L50 (5–8), and largest contig (3.6–6.1 Mb). Assembly completeness was evaluated with BUSCO v5.2.2 (38) using the fungi_odb10 and eukaryota_odb10 datasets, reporting completeness scores of 88%–89% and 94%–95%, respectively. To assess chromosome completeness, the assemblies were analyzed for telomeric sequences using find_telomeres.py v0.2 (https://github.com/markhilt/genome_analysis_tools), identifying between 5–8 telomere-to-telomere contigs per genome.

### Read mapping and coverage analysis

CCS reads were mapped to the assembled genomes using minimap2 v2.28 (39) with the parameters -ax map-hifi. Mapping rates (≥99.8%) and sequencing depth (range: 217–411) were then calculated using Samtools v1.14 (40). Mapping results were visualized in Jalview v2.11.4.1 (41).

### Target locus identification and validation

Target recombination loci were identified using BLAST v2.15.0 (42). tblastn searches were performed with GAPDH, Myo17D, and URA3 sequences to locate genomic integration sites, and additional searches were conducted for hph, mRuby3, and AmpR (from plasmid backbone) to detect off-target integrations. The same process was repeated for each haplotype assembly. Integration sites were validated using blastn searches against raw HiFi reads, applying an e-value threshold of 0.0001 and sequence identity >90%.

### Protein extraction and immunoblot analysis

To extract total protein, we centrifuged zoospores harvested from two 4-day-old agar plates at 2500 × g for 5 minutes. We then resuspended the pellet in 80 µl lysis buffer (final concentration: 62.5 mM Tris pH 6.8, 70mM SDS, 10% (v/v) glycerol, 5 mM EDTA, 1X HALT protease inhibitor, 1mM PMSF). After 5 minute incubation at 95 °C, we centrifuged samples for 3 minutes at 20,000 g. We quantified protein concentration in samples using a DC protein assay kit (Bio-Rad, Cat. No. 5000112) and diluted samples to 2 µg/µl in lysis buffer. For immunoblotting, samples were diluted to 1 µg/µl in SDS sample buffer (final concentration: 60 mM Tris, 2.5% SDS, 0.025% bromophenol blue, 10% (v/v) glycerol, 0.1 M DTT) and 10 µl of each sample were loaded into wells of a 4-20% polyacrylamide gel (Bio-Rad, Cat. No. 4568096). After gel electrophoresis, we transferred the gel overnight onto a PVDF membrane (Cytiva, Cat. No. 10600102) in Towbins buffer (25 mM Tris, 192 mM glycine, 20% (v/v) methanol). We blocked the membrane in TBS-T (20 mM Tris pH 7.5, 137 mM NaCl, 0.5% (v/v) Tween-20) + 3% skim milk (Difco) for 1 hour, then probed with either rabbit anti-GAPDH primary antibody (Abcam, Cat. No. ab22555) at 1:800 dilution in TBS-T + 1% skim milk or rabbit anti-TagRFP primary antibody (ThermoFisher Scientific, Cat. No. R10367) at 1:1,000 dilution in TBS-T + 1% skim milk for 2 hours. We then incubated the membrane with HRP-conjugated anti-Rabbit IgG secondary antibody (ThermoFisher Scientific, Cat. No. G-21234) at 1:10,000 dilution in TBS-T + 1% skim milk for 1 hour. Hybridization signals were visualized with ECL Western Blotting Detection System (Cytiva, Cat. No. RPN2232) and a G:Box XX9 gel imager.

### Virulence testing

All procedures involving live frogs in this study were conducted at the University of Michigan Animal Care Facilities following protocols approved by the University’s Institutional Animal Care and Use Committee (IACUC protocol PRO00011269). Animals were cared for and housed according to previously established methods (43). Briefly, during the experiments, adult *Hymenochirus boettgeri* were housed individually in 1 l tanks filled with reverse osmosis (RO) water, with conductivity adjusted to 900 ± 100 µS/cm. Each tank was enriched with sterilized gravel (greater than 1 cm in diameter) and included one sterilized PVC pipe. The tanks were covered with plastic lids that featured small holes to promote air circulation. Animals were monitored daily and fed a diet of small-granule fish food (BUG BITES Tropical Formula, Mansfield, MA, USA) and gamma-irradiated bloodworms (*Chironomus* sp.; Tropical Marine Center, UK) ad libitum three times a week. To remove nitrogenous residues, water changes were performed five hours after feeding. All experiments conducted in this paper with *H. boettgeri* were carried out in a temperature-controlled room set at 20°C ± 0.5°C, with a 13:11 light–dark cycle.

For the virulence experiment, JEL423 and *Bd*-EK42-3 were grown on 1% tryptone agar plates for 3 days at 20°C. Hygromycin B was not included in the media to grow *Bd*-EK42-3, as this could potentially affect the animals’ microbiome. However, *Bd*-EK42-3 was cultured on selective plates and transferred to non-selective plates for only 3 days. Additionally, cultures of *Bd*-EK42-3 on 1% tryptone agar plates were checked before the experiment, and no changes in fluorescence were detected. To obtain zoospores for inoculum preparation, we flooded plates of both strains and *Bd*-free 1% tryptone agar plates (used for the sham inoculation) with 2 ml of sterile water and incubated them at room temperature for one hour to stimulate zoospore release. The liquid was then harvested, filtered through a 40 µm mesh filter to remove large clumps of sporangia (Fisher Scientific, Cat. No. 22-363-547), and the cells were diluted 1:1 with trypan blue (Gibco, CAT 15250061). Cell counts were performed using a hemocytometer. The filtered cells were then diluted to 1 × 10⁵ cells/ml in 5 ml of sterile pre-conditioned RO water.

We selected *H. boettgeri* adults of approximately the same size (mean SVL = 2.74 ± 0.19 cm, n = 24) and weight (mean = 1.32 ± 0.28 g, n = 24) and randomly assigned them to one of two treatment groups or a control group, ensuring a balanced sex ratio (i.e., 4 females and 4 males). Prior to the experiment, all animals were swabbed on the skin following the protocol of Hyatt et al. (44) using disposable gloves, and all tested negative for *Bd* (see *Bd* load quantification section below).

For pathogen exposure, we placed animals individually into 50 ml conical tubes containing 5 ml of either the inoculum or sham inoculum. The tubes were capped, and the frogs were left to sit for approximately 3.5 hours, with periodic checks to ensure they remained submerged in the water and were not resting on the sides of the tube. After inoculation, the animals were placed individually in 1 l tanks, including the inoculum they were exposed to, and monitored three times daily for signs of disease (*e.g*., lethargy, lack of appetite, skin shedding, and lack of righting response). Upon displaying signs of advanced disease or at the end of the experiment, all animals were swabbed and euthanized using tricaine methanesulfonate (MS-222; 0.5 g/l; Syndel, Ferndale, WA, USA) followed by double pithing. The date of death was recorded and swabs were stored at -20°C before DNA extraction.

We extracted *Bd* DNA from skin swabs using PrepMan (Applied Biosystems CAT 4318930), following the manufacturer’s protocol. The extracted DNA was subsequently used as input for a standard quantitative PCR (qPCR) assay (45), performed on a QuantStudio 3 (Applied Biosystems, ThermoFisher Scientific). The mean *Bd* loads in zoospore equivalents (ZE) were then calculated from standard curves generated using DNA extractions from seven serial dilutions (ranging from 10⁶ to 10⁰ ZE) of JEL 423. Samples were run in duplicate to ensure accuracy, and only those samples that amplified ≥1 ZE in both replicates were considered positive for *Bd*. In cases of inconsistent results, repeat testing was performed, and samples were considered positive when two out of three replicates were positive. The mean *Bd* load in ZE was recorded for each positive sample. Negative controls were included in all qPCR assays to account for potential contamination.

Survival curves were generated using the Kaplan-Meier method, implemented via the *survfit* function from the *survival* package (46). We compared the survival curves between the two treatments using log-rank tests through the *survdiff* function from the same package. Additionally, to compare the *Bd* load (log-transformed) at the time of death between individuals exposed to the *Bd* JEL423 and *Bd-*EK42-3 strains, we performed an analysis of variance (ANOVA) using the *aov* function from the *stats* package (42). All analyses were conducted in R version 4.0.2 (42).

### Live frog and webbing explant imaging

To visualize *Bd* on live animals, we exposed three individuals of *H. boettgeri*, each with either 5 × 10^5^ zoospores of wild type JEL423 or 5 × 10^5^ zoospores of the GAPDH-mRuby3 expressing stain (*Bd*-EK42-3). All individuals were approximately the same size (∼30 mm) and weight (∼1 g). We maintained each *Bd* strain as described above. On day five or seven of infection, animals were anesthetized using 0.5 g/l MS-222 for at least 15 minutes. We placed the sleeping frogs onto 48 x 60mm #1 glass coverslips and ensured their hands and feet were spread out while keeping the legs as flat as possible. We imaged whole animals on an inverted microscope (DMi8 with THUNDER Imager; Leica) with a 10X, 0.3 NA air objective using LAS X software (v3.9.0.28093). Images were taken using widefield fluorescence microscopy with an excitation wavelength of 555 nm to visualize GAPDH-mRuby3. Images of both the dorsal and ventral sides of each frog were taken at room temperature.

After imaging, the animals were returned to 0.5 g/l MS-222 for a minimum of 10 minutes, then euthanized by double pithing. We then used a scalpel to remove webbing from the hind legs and mounted the skin explants in water on slides using 22 x 22 mm #1 coverglass, ensuring the webbing was laid as thin and flat as possible. Slides were sealed with clear nail polish. We imaged the slides right away at high resolution on an upright microscope (Examiner Z1; Zeiss) with a 63X, 1.4 NA oil objective using Zen software. Images were taken using scanning point confocal fluorescence microscopy (LSM880, Zeiss) with an excitation wavelength of 561 nm and emission wavelength range between 579 and 695 nm to visualize GAPDH-mRuby3 and transmitted light of the 561 nm excitation wavelength to view whole cells. We took z-stack images to encompass whole *Bd* cells in the frog epithelium. All images were taken at room temperature.

### Data, materials, and software availability

Whole genome sequences are available through NCBI Sequence Read Archive BioProject: PRJNA1311079. Deposited strains are available for request from the CZEUM repository (https://czeum.herb.lsa.umich.edu/).

## ACKNOWLEDGEMENTS

We thank Sam Lord for comments on the manuscript, James Chambers (Director, University of Massachusetts Light Microscopy Facility), Gregg Sobocinski (Managing Director, University of Michigan Department of Molecular, Cell, and Developmental Biology Microscope Imaging and Cell Analysis Core Facility), and Ravi Ranjan (Director, University of Massachusetts Genomics Resources Laboratory) for technical support and Edgar Medina for sharing plasmid constructs. This work was funded by the Gordon and Betty Moore Foundation grant (award #9337) to L.K.F.-L. and T.Y.J., who are Canadian Institute for Advanced Research (CIFAR) fellows in the Fungal Kingdom: Threats and Opportunities program.

---

**Figure S1:**
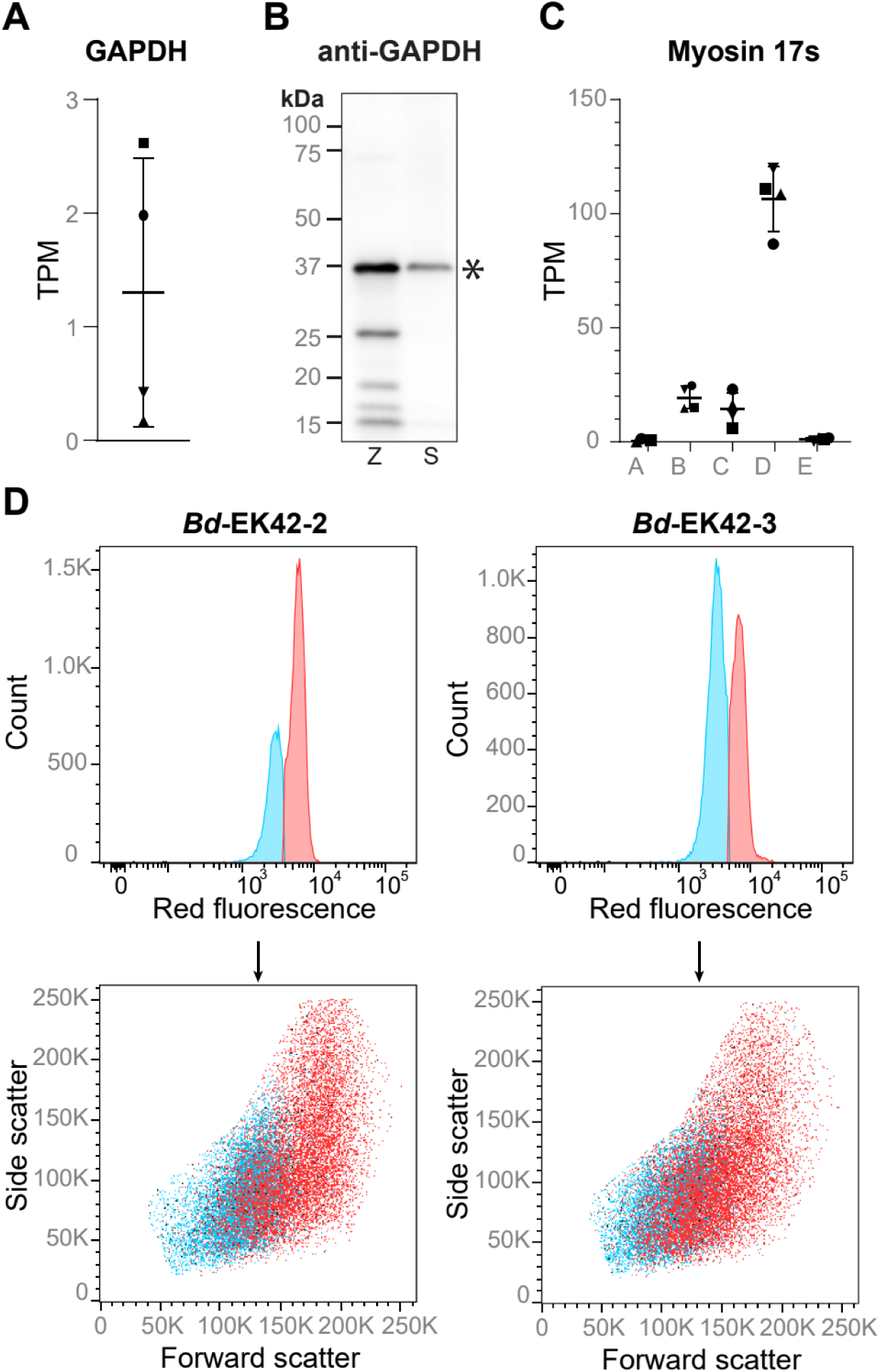
Additional information about *Bd* gene expression and fluorescence of transformed cell lines. **(A)** Transcripts per million (TPM) of GAPDH in WT JEL423 zoospores. **(B)** Immunoblot analysis of GAPDH expression in WT JEL423 zoospores (Z) and 24 hour sporangia (S). Star denotes expected size band size for native GAPDH protein (36 kDa). **(C)** TPM of the 5 Myosin 17s in WT JEL423 zoospores. In the current *Bd* JEL423 genome, Myo17B is split across two gene annotations, separating the motor domain from the chitin synthase domain. Here, Myo17B TPM was based on the myosin motor domain annotation alone. **(D)** Backgating of putative subpopulations GAPDH-mRuby3 cells onto forward scatter vs. side scatter shows the two distinct fluorescence peaks emerged from differently sized subpopulations. Scatter plots show single cells only, after gating for cells and doublet discrimination: mRuby3 negative cells shown in black, weakly fluorescent in blue, strongly fluorescent in red.

**Figure S2:**
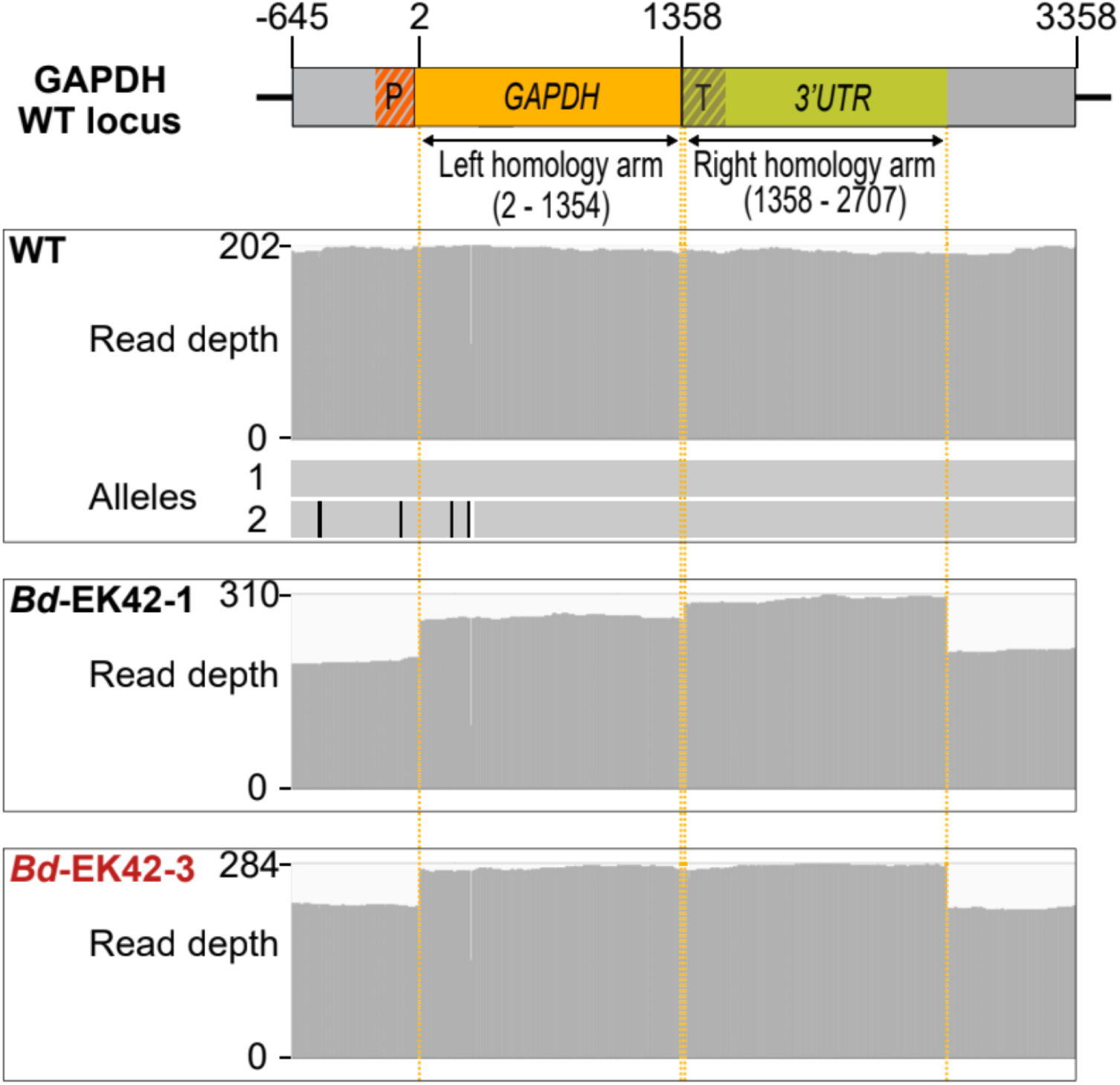
GAPDH wild type (WT) locus structure and read depths according to whole genome assemblies. Haplotype resolution was achieved through SNPs (black lines) and a 3 bp deletion (white line) specific to allele 2. Read depth shows the number of reads that map to wild type (WT) and edited loci. The transgenic clones show higher read depth in the region with sequence identity to the homology arms, supporting the presence of additional copies of the homology arm sequences in these strains.

**Figure S3:**
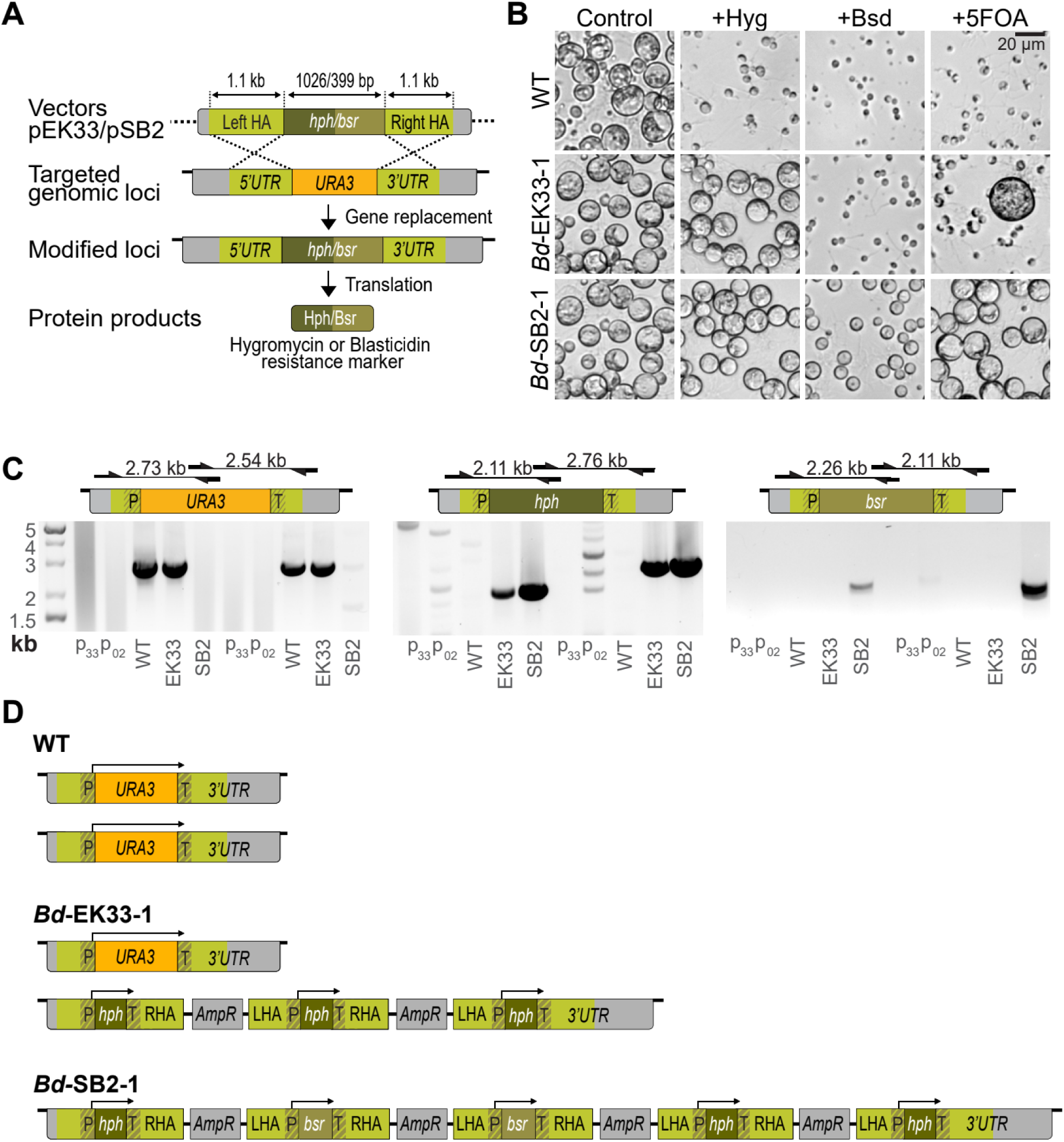
Gene replacement of *Bd URA3* by sequential transformation with two selection constructs: **(A)** Schematic representation of the experimental strategy used to generate stable *BdURA*3 knock-out through sequential gene replacement using homologous recombination. Cells were sequentially transformed with the vector pEK33 then pSB02 (*hph:* hygromycin resistance gene, bsr: blasticidin resistance gene, HA: homology arm). **(B)** Bright field images of wild type (WT), *Bd*-EK33-1, and *Bd*-SB02-1 lines cells grown in media supplemented with hygromycin B, blasticidin S or 5-FOA for 3 days. **(C)** (Top) Diagrams showing primer locations and amplicon sizes of different primer pairs to native *URA3* locus or transformed loci. (Bottom) Gel electrophoresis of genotyping PCR confirming presence of the native *URA3* locus and *hph* or *bsr* locus specific integration into WT, *Bd*-EK33-1 (EK33), and *Bd*-SB02-1(SB2) lines (P_33_ = pEK33 plasmid control, P_02_=pSB02 plasmid control). **(D)** Diagram of reconstructed alleles of the *URA3* loci of WT, *Bd*-EK33-1 and *Bd*-SB02-1 based on whole genome sequencing assembly results. Note: WGS of *Bd*-SB02-1 recovered only a single allele of the *URA3* locus, but read depth analysis indicates that there are likely two copies of this allele present in the genome. The arrows above each allele denote predicted open reading frames (P: promoter, T: terminator, *hph*: hygromycin resistance gene, *bsr*: blasticidin resistance gene, L/RHA: left/right homology arm).

**Table S1:** Overview of electroporation-based stable *Bd* transformation. A general timeline outlining the various steps involved in generating transgenic *Bd* lines using gene targeting vectors.

| Day | Procedure | Details |
| --- | --- | --- |
| 1 | Electroporate cells |  |
| 1-5 | Recover electroporated cells in non-selective liquid media |  |
| 2 | Inoculate antibiotics selection test | Inoculate 1 out of 7 wells for each sample |
| 5 | Assess antibiotics containing well for transient growth | Check for growing sporangia and release of motile zoospores |
| 5 | Plate samples onto selection and control agar plates | One of each type of plate per sample |
| 9-21 | Check selection plates for transient transformants and control plates for cell survival |  |
| 23-26 | Inspect selection plates for stable growth (>100 swimming zoospores per field of view) | Required incubation duration varies between transformation trials and gene targeting vectors |
| 26-50 | Single colony isolation | Two consecutive rounds |
| 50-55 | Culture amplification |  |
| From 92 | Cryopreservation and downstream analysis |  |

**Table S2:**
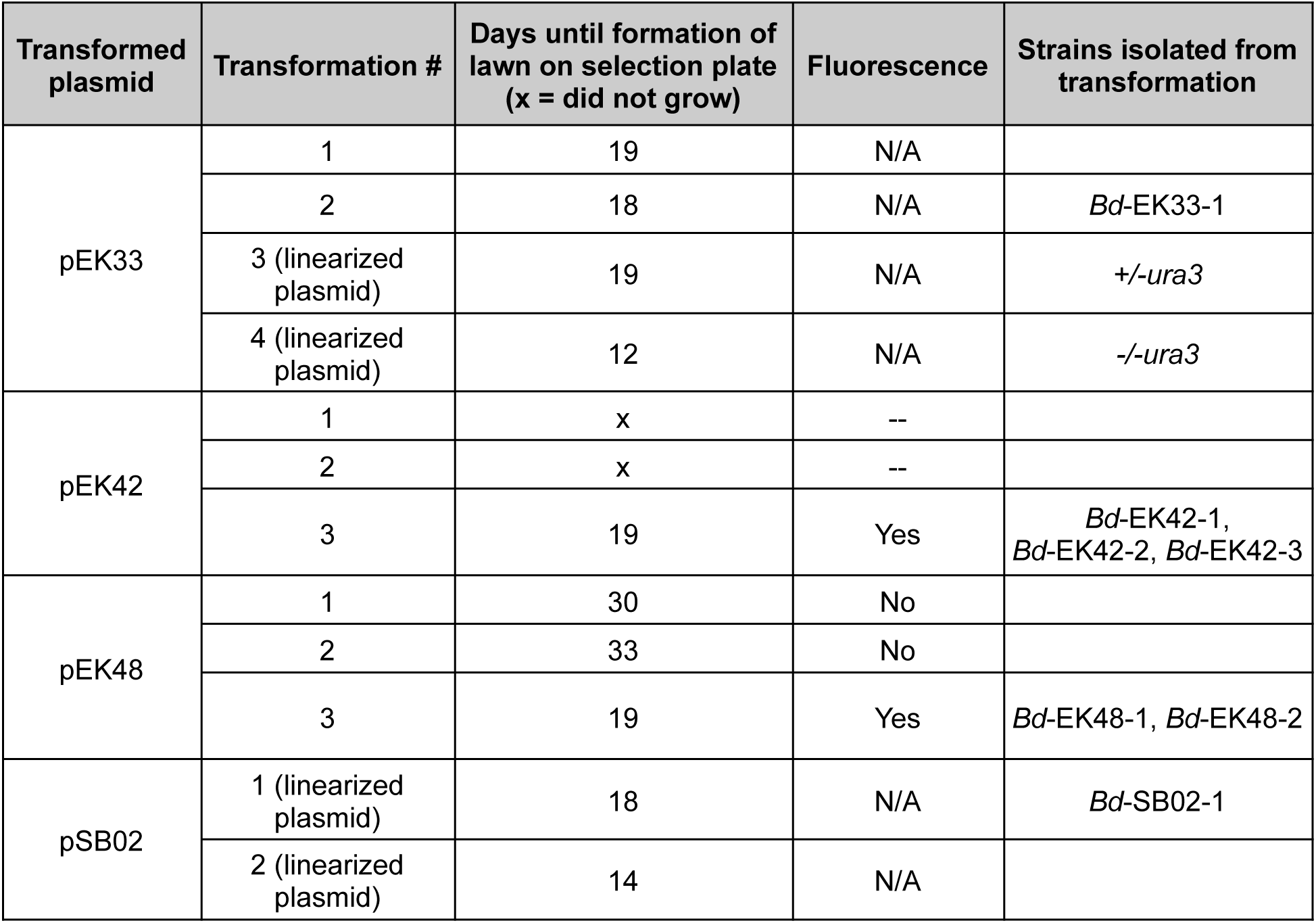
Statistics on transformation success. Statistics for the generation of stable fluorescent transformants of *Bd* JEL423 using gene targeting vectors.

**Table S3:**
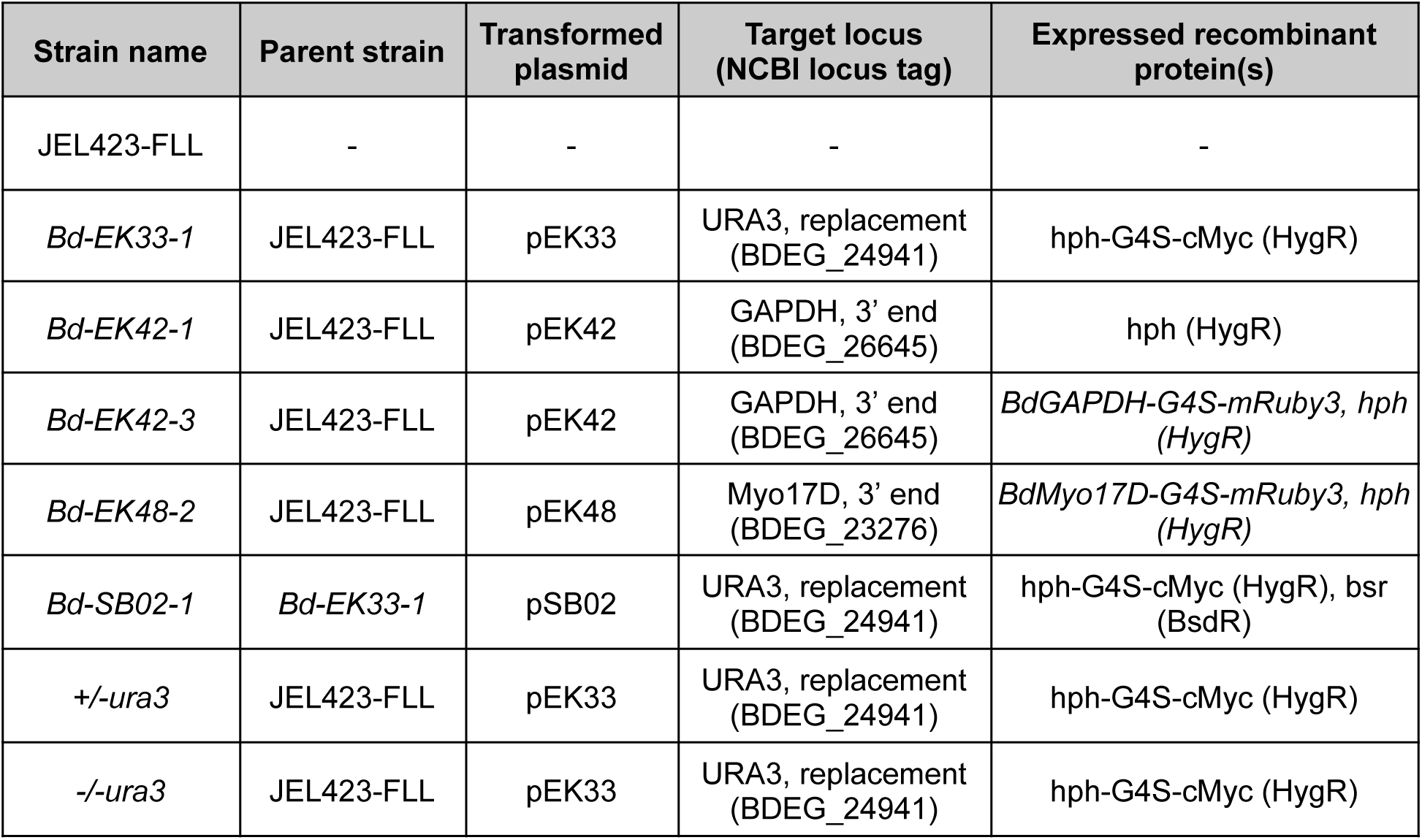
*Bd* strains available from the Collection of Zoosporic Eufungi at the University of Michigan (CZEUM). Accessible at czeum.herb.lsa.umich.edu.

## SUPPLEMENTARY VIDEO AND DATA SET LEGENDS

**Dataset S1: *Batrachochytrium dendrobatidis* gene targeting vectors used in this study.** This data set lists the names, source, and cloning strategy used for each plasmid. The JEL423 genome assembly (GCA_000149865.1) was used to construct gene models.

**Dataset S2: Primers used in PCR screenings.** Dataset containing primers used in this study.

**Dataset S3: Assemblies of relevant loci.** Sequences of assembled loci relevant to this study.

**Movie S1: Video of Myo17D-mRuby3 localization during mucin-induced encystation shown in Fig. 4G**. Images were taken every 5 seconds in DIC and mRuby3. Time is displayed in minutes:seconds.

